# CH_4_ transport in wetland plants under controlled environmental conditions – untangling the impacts of phenology

**DOI:** 10.1101/2023.10.08.561392

**Authors:** Mengyu Ge, Aino Korrensalo, Anuliina Putkinen, Raija Laiho, Lukas Kohl, Mari Pihlatie, Annalea Lohila, Päivi Makiranta, Henri Siljanen, Eeva-Stiina Tuittila, Markku Koskinen

## Abstract

- Methane (CH_4_) fluxes at plant surfaces are the net result of transport of soil-produced CH_4_ and within-plant CH_4_ production and consumption, yet factors and processes controlling these fluxes remain unclear.
- We conducted high-frequency automated CH_4_ flux measurements from shoots of *Carex rostrata* (sedge), *Menyanthes trifoliata* (forb) and shrubs (*Betula nana*, *Salix lapponum*) during early, high and late summer in a climate-controlled environment to assess the effects of environmental variables, seasonality and CH4 cycling microbes in the CH4 flux. Measurements were conducted from intact plant-soil samples collected throughout growing seasons 2020 and 2021 from Lompolojänkkäfen, northern Finland.
- All studied species showed seasonal variability in CH_4_ fluxes. The CH4 fluxes were not impacted by light level, while out of the studied species, porewater CH_4_ concentration increased fluxes from all but B. nana. Air temperature only and negatively affected CH4 flux from C. rostrata. Both methanogens and methanotrophs were detected in aboveground parts of *S. lapponum* and *M. trifoliata*, methanotrophs in *B. nana*, while neither were detected in *C. rostrata*.
- Our study demonstrates that the seasonal phase of the plants regulates CH4 flux they mediate across species, which was not observed in the field. The detection of methanogens and methanotrophs in herbs and shrubs suggests that microbial processes may contribute to their CH_4_ flux.

## Introduction

Methane (CH_4_) is a powerful greenhouse gas with a global warming potential 34 times larger than carbon dioxide over 100 years (Saunois *et al*., 2016). Undrained peatlands are an important source of CH_4_, accounting for *c.* 12.2% of global CH_4_ emissions (Rydin & Jeglum, 2006). These emissions are enabled by the high water-table level (WTL) which is a pre-requisite for many peatland plant species to thrive, and can be both stimulated and attenuated by members of the plant community. Vascular plants can stimulate CH4 emissions by offering substrates for microbial CH_4_ production and by transporting this CH_4_ to the atmosphere (Greenup, 2000; Joabsson & Christensen, 2001; Ström *et al*., 2003; Barba et al., 2019). They can attenuate CH4 emission by transporting oxygen from shoots to roots, thereby stimulating CH_4_ oxidation in the soil (Heilman and Carlton, 2001; Robroek et al., 2015). The net effect of the plant community on the CH4 balance of a peatland depends on the interplay between these stimulating and attenuating tendencies, the spatiotemporal and interspecific variation of which is still not well known.

Plant-mediated CH_4_ transport allows soil-produced CH_4_ to bypass oxidation in oxic surface peat and thus is a crucial mechanism increasing the total ecosystem CH_4_ emissions both in open wetlands and woody ecosystems (Shannon *et al*., 1996; Pangala *et al*., 2017; Barba *et al*., 2019; Ge *et al*., 2023; Machacova *et al*., 2023). Environmental factors such as soil water table level (WTL), temperature and porewater CH_4_ concentration ([CH_4_]_pw_) have been regarded as important controls on ecosystem CH_4_ emissions from wetlands (Joabsson & Christensen, 2001; Lai *et al*., 2014a). Yet, the effects of these controls on plant-mediated CH_4_ transport remain unclear with contrasting results even for the same species. For example, Noyce *et al*. (2014) reported that increasing WTL lead to high CH_4_ emissions by *Carex rostrata*, while Ge *et al*., (2023) observed no such correlation with the same species. Similarly, [CH_4_]_pw_ has been found to positively influence plant-mediated CH_4_ emission (Aulakh *et al*., 2000; Ding *et al*., 2005), while no such effect was observed in our earlier study where plant-mediated CH_4_ flux (fCH4p) was measured from common boreal peatland plants with different morphological traits (Ge *et al*. (2023)).

Recent studies suggest that environmental controls are less important than species-level differences (Korrensalo *et al.,* 2021) and seasonality (Ge *et al*., 2023) in determining plant-mediated CH4 transport. It seems evident that species-specific plant morphology and phenology (the development and senescence over the growing season) can affect the CH4 transport. Nevertheless, it remains unclear which plant parts and properties determine the resistance to CH4 transport and release, even for the most well-studied genus, *Carex spp.*. Studies based on clipping plant aboveground parts either suggest that the plant shoots do not restrict CH4 transport (Nouchi *et al*., 1990; Kelker & Chanton, 1997; MacDonald *et al*., 1998; Kutzbach *et al*., 2004; Schimel, 1995), or opposingly, that they strongly control the CH4 release (Morrissey *et al*., 1993; Schimel, 1995; Garnet *et al*., 2005). Phenology may affect CH_4_ transport through changes in plant biomass (Koelbener *et al*., 2010), root length (Noyce, 2009) and permeability (Henneberg *et al*., 2012), aerenchyma size (Fagerstedt, 1992), stomatal conductance (Morrissey *et al*., 1993), and stimulation of rhizospheric CH_4_ production (Lai *et al*., 2014b) and oxidation (Kankaala *et al*., 2005). The seasonal variability of plant-mediated CH4 flux may also differ between plant species (Ge et al., 2023). However, most previous studies have been conducted in the field, where abiotic conditions typically correlate with phenological changes in plants. Studying plant-mediated CH4 flux under controlled environmental conditions would allow investigating the relative importance of both the species identity and species-specific seasonality in regulating the flux.

Plants are not merely transporting soil-derived CH4 to the atmosphere but may also host CH4 producing and consuming microbes (Larmola *et al*., 2010; Putkinen *et al*., 2021). These processes, however, are rarely accounted for in ecosystem CH4 flux studies even though recent studies indicate that both microbial CH4 production and consumption (Jeffrey *et al*., 2019; Jeffrey *et al*., 2021; Putkinen *et al*., 2021). While the microbiomes of both herbaceous and shrub species remain poorly examined, the recent detection of CH_4_ cycling microbes in tree leaves (Putkinen *et al*., 2021) and stems (Feng *et al*., 2022), and the well-known association between *Sphagnum* mosses and methanotrophs (Larmola *et al*., 2010; Putkinen *et al*., 2014), support their wider evaluation within other plants, especially those adapted to CH_4_-rich environments like peatlands.

In this study, we investigate the seasonality and species-specific differences in CH_4_ flux of shoots of common boreal peatland herbs (sedge *C. rostrata* and forb *M. trifoliata*) and shrubs (*B. nana* and *S. lapponum*) in controlled environmental conditions. The aims were to (i) evaluate how the phase of the growing season and abiotic factors (temperature, [CH_4_]_pw_ and PAR) interact in controlling plant-mediated CH_4_ flux; (ii) identify which plant part is the main location of CH_4_ release from *C. rostrata*, one of the most abundant sedge species in northern peatlands; and (iii) determine whether plant shoots host microbes that produce or oxidize CH4. To disentangle the effects of season mediated through changes in environmental conditions and plant phenology, we collected plant-soil mesocosms containing the target species throughout growing seasons in 2020 and 2021 and placed them in a climate-controlled cabinet for automated measurements under uniform environmental conditions (Kohl et al., 2021). To reveal whether leaves of C. rostrata restrict the transport of CH_4_, a foliage clipping experiment was conducted. To evaluate CH4 production and consumption processes occurring inside of plant shoots, a ^13^C-labelling measurement was conducted, and the presence and diversity of methanogens and methanotrophs within plant shoots were measured. We hypothesized that i) phenological stage of the plants modifies the fCH_4p_, ii) the fCH_4p_ of all investigated species increases with PAR, temperature and [CH_4_]_pw_, iii) leaves are the main location for CH_4_ release from *C. rostrata*, and iv) CH4-related processes occurring inside plant shoots include plant-derived and microbial CH4 production and oxidation.

## Materials and methods

### Plant sampling and growth condition

Mesocosms of *Carex rostrata*, *Menyanthes trifoliata*, *Betula nana* and *Salix lapponum* were collected at Lompolojänkkä(67.99 °N, 24.21°E), a northern boreal fen (Ge *et al*., 2023) located in Finnish Lapland. The plants were collected in growing seasons 2020 (*C. rostrata* and *B. nana*) and 2021 (*M. trifoliata* and *S. lapponum*). To investigate the impact of phenology on plant CH4 transport, we collected the mesocosms of each species at three time points, which represent phenologically distinct phases. The sampling times were chosen based on earlier studies at the site (Ding *et al*., 2003; Zhang *et al*., 2020; Ge *et al*., 2023); early summer leaf out (early-mid June), high summer during the leaf area maximum (July–early August) and autumn senescence (mid-September).

There were three mesocosms per species per campaign. The total number of mesocosms throughout the observations was less than nine due to system malfunctions and the quality control of closure data (Table S1). The plant individuals were the same ones measured in Ge et al. (2023), enabling the comparison of the results obtained in situ and in the climate-controlled measurements. To reduce the disturbance to the plants, each mesocosm included the peat surrounding the roots of the target plant individual and was placed in a plastic bucket (10 L). We dug the peat up to 40 cm depth to avoid disturbance to the roots, of which the majority are located above the depth of 30 cm in boreal sedge fens (Mäkiranta *et al*., 2018). All mesocosms passed careful visual inspection of the success in preserving root systems, with no shrub and herb mesocosms having the end of the rhizome and/or coarse roots sticking out of the sampled peat.

The mesocosms were stored in an environment-controlled room at 4 °C with a 16-h photoperiod before being transported to the climate-controlled cabinet to ensure uniform growing conditions and to prevent the individuals from progressing to different phenological stages before measurements. To keep the peat water-saturated, the mesocosms were watered with water collected from the hole left after digging out the specimen and stored in an airtight bottle stored in the same environment-controlled room. The size of the climate-controlled cabinet limited measurements to two mesocosms at a time. We therefore conducted simultaneous measurement of two mesocosms of different species, with the remaining two mesocosms of each target species stored for one or two weeks, respectively, in the environment-controlled room.

### Porewater CH_4_ concentration

Before starting the measurements, a silicone tube attached to a polytetrafluoroethylene (PTFE) tube was placed at the bottom of each bucket (Kammann *et al*. (2001). The outer diameter and wall thickness were 16 and 3 mm for the silicone tube, and 6 mm and 1 mm for the PTFE tube, respectively. The top end of the PTFE tube was sealed with a 3-way stopper through which gas samples were extracted by a syringe (20 ml) and transferred into pre-evacuated vials (12 ml, Labco Limited, UK) daily. CH_4_ concentrations were analysed by a gas chromatograph (GC, Agilent 7890A). The measured gas phase concentrations (ppm CH_4_) inside the tube were then converted to concentrations (µmol CH_4_ l^-1^) in the surrounding porewater based on Henry’s law:

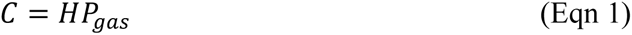

Where C is the solubility of a gas at a fixed temperature in a particular solvent (mol l^-1^), here CH_4_ concentration in porewater; P_gas_ is the partial pressure of CH_4_ (bar); and H is Henry’s law constant (mol l^-1^ bar^-1^) which for CH_4_ at 298K is 0.0014 mol l^-1^ bar^-1^.

### Experimental set-up

The measurements were conducted in a climate-controlled cabinet (Fitoclima 1200, Aralab, Portugal, Fig. 1), where PAR, temperature and the relative humidity of the air (RH) could be regulated. Each measurement cycle lasted approximately one week and was conducted with two mesocosms hosting two different species at a time. Throughout the observations, regular light cycles (18 h light, 6 h darkness) were applied and RH was held at 50% (Table 1). The PAR was kept at 600 and 100 µmol m^-2^ s^-1^ intensity for the daytime and nighttime, respectively. We did not set PAR to zero as the field site also had light at night over most of the growing season. Temperature was kept at 15 °C for two days and then increased to 20 °C.

**Fig. 1.**
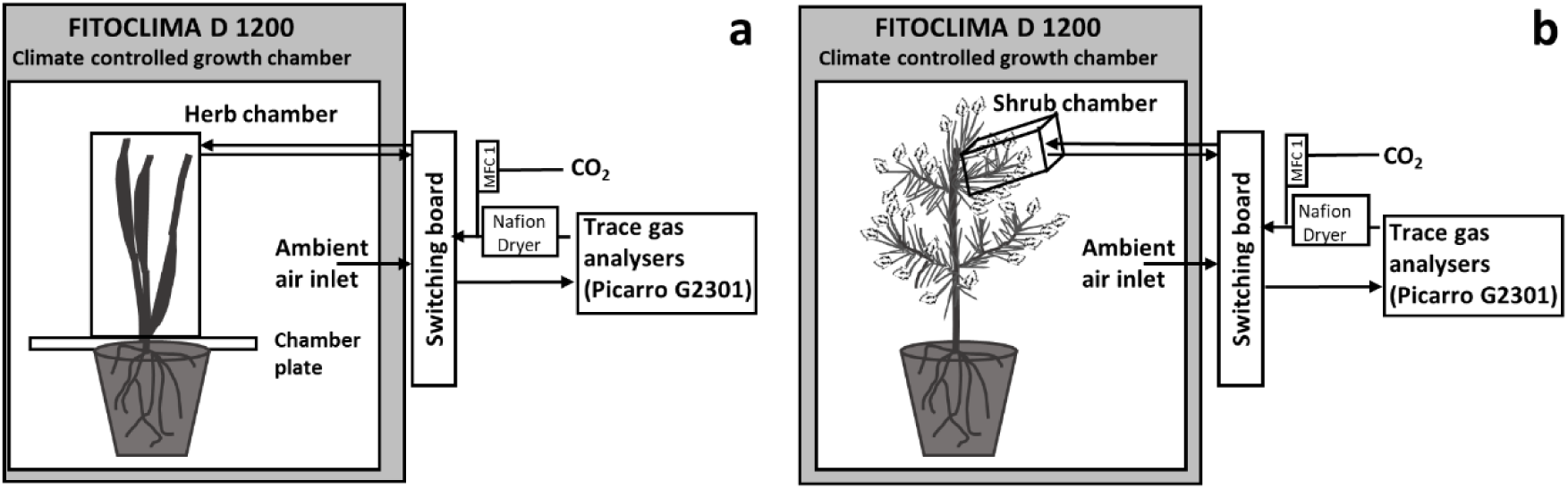
The controlled-environment chamber system, consisting of a climate-controlled growth chamber, and two different measurement chambers for observing CH_4_ fluxes from herbaceous species *Carex rostrata* and *Menyanthes trifoliata* (a, herb chamber), and from shrubs *Betula nana* and *Salix lapponum* (b, shrub chamber). Figure modified after Kohl *et al*. (2021).

**Table 1.** Climate-controlled cabinet settings and treatment protocol. The foliage clipping treatment conducted only on *C. rostrata*.

| Setting | Day 1 | Day 2 | Day 3 | Day 4 | Day 5 | Day 6 | Day 7 |
| --- | --- | --- | --- | --- | --- | --- | --- |
| RH (%) | 50 | 50 | 50 | 50 | 50 | 50 | 50 |
| T (°C) | 15 | 15 | 20 | 20 | 20 | 20 | 20 |
| Light / Dark (h) | 18/6 | 18/6 | 18/6 | 18/6 | 18/6 | 18/6 | 18/6 |
| Clipping foliage of *C. rostrata* | no | no | no | no | yes | yes | yes |

The clipping treatment was conducted on C. rostrata to reveal whether the CH4 transport of the plant was restrained by passage through the leaves. Specifically, the foliage of C. rostrata was removed and only the stem was left. The clipping treatment was conducted on the fifth day of the measurement cycle, during the 20 °C air temperature phase, and measurements continued for 3 days after the treatment (Table 1). We only conducted the clipping treatment on *C. rostrata* as it is one of the most important and frequent species in northern peatlands and also has been found to be an efficient CH4 emitter (Noyce *et al*., 2014; Ge *et al*., 2023). Two mesocosms per campaign were treated by clipping. The effect of clipping was assessed by comparing the measurements made during the 20 °C air temperature phase before and after the treatment.

### CH_4_ flux measurements

Due to different growth forms of herbs and shrubs, two different measurement chamber designs were used for measuring CH_4_ fluxes from their shoots (Fig. 1). We used the chamber design developed by Korrensalo *et al*. (2021) for the herbs, hereafter called “herb chamber”. It consisted of two plexiglass plates and a transparent polymethyl methacrylate (PMMA) chamber (4 L). Measurement of shoot fluxes separate from the rest of the environment was achieved by passing the specimen through the gap between the two plexiglass plates being held together by a hinge and then covering it with the chamber.

Unlike herbs, we measured CH_4_ fluxes from parts of shrub shoots by using a 1 L chamber system, hereafter called “shrub chamber” (Toivo Pohja, Juupajoki, Finland). It consisted of an aluminium bottom and back plate, and a transparent PMMA cover. The measured stem was put through a notch in the back plate and sealed with adhesive putty (Blu Tack, Bostik SA).

CH_4_ fluxes were measured in 10-min long closures and there were 72 closures per species per day. During each closure, sample gas in the headspace was circulated in a closed loop between the analyser (Picarro G2301 and G2201i) and the chamber. After each closure, the chamber was flushed with air taken from the cabinet for 10 minutes while the other chamber was under measurement. We conducted empty chamber measurements to calculate background fluxes after finishing all campaigns in 2020. Chamber tightness was monitored with daily open-cabinet measurements. Opening the cabinet doors lowered CH_4_ concentrations within the cabinet, which would have had effects on the apparent CH_4_ flux if there was significant leak in the chamber. This was not observed as fluxes measured with the cabinet doors open did not differ from those measured with the doors closed.

CH_4_ fluxes through plants and the empty chamber were calculated using the least squares methods, defining the best linear fit for the change of concentration over time:

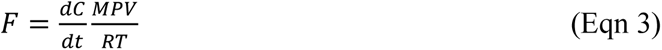

where dC/dt is the change in CH_4_ (ppm) concentration over time (s), M is the molar mass of measured gas (16042 mg for CH_4_), P is the atmospheric pressure (101325 Pa), V is the volume of the measurement system (m^3^), R is the molar gas constant (8.3144598 J K^-1^ mol^-1^), and T is the chamber temperature (K).

We used a custom R script for quality control of each closure. Closures with nonlinear changes in CH4 were identified to be caused by chamber leak and were excluded. We also discarded closures conducted during system malfunctions, or opening of the cabinet door to dilute cabinet CH4 concentration. In total, 25% of the closures were rejected. The mean background flux of the empty herb and shrub chambers were very small (-2.08×10^-6^ and - 1.41×10^-6^ mg d^-1^, respectively).

After each campaign, we measured plant surface area (leaf area and stem area for herbs and shrubs, respectively) enclosed in the chambers. The single-sided leaf area was measured by a digital scanner and ImageJ software (Ferreira & Rasband, 2012). A digital caliper meter (1150D; Bahco, Eskilstuna, Sweden) was used to measure the stem diameter of shrubs at the point of entry to the shrub chamber, from which we calculated the cross-sectional stem area. After that, CH4 flux from plants was divided by plant surface area to estimate the area-specific fCH4p. We use the stem area instead of leaf area for shrubs because in the autumn campaign, senescence had set in and leaf area was zero and we wanted to use the same parameter for normalizing the flux of the shrubs in all campaigns.

### 13C labelling

While aerenchymatic herbs have long been regarded as efficient channels for soil-produced CH4, with shrubs the main source of CH4 emitted through themis poorly studied. We therefore conducted a labelling experiment on *S. lapponum* to reveal whether the CH_4_ emitted by its shoots originated from the peat following an approach previously used to confirm peat as the source of CH4 emitted by *B. nana* (Koskinen *et al*., 2021). Two ml ^13^C-labelled CH_4_ (99% ^13^C, Merck 490229) were mixed with 28 ml dinitrogen (N_2_) gas and injected into the silicone tube installed at the bottom of the mesocosms for porewater CH_4_ concentration ([CH_4_]_pw_) measurements. All three mesocosms of S. lapponum from each campaign were labelled once on the fifth measurement day and ^13^CH_4_ emissions from shoots were recorded until the end of the measurement cycle. The δ^13^C-CH_4_ of shoot-emitted methane was calculated through Keeling plots, i.e., by plotting the measured δ^13^C-CH_4_ over the inverse concentration and estimating the intercept of a linear fit in these plots (Pataki *et al*., 2003). Similarly, we estimated the initial soil δ^13^C-CH_4_ based on CH_4_ emitted from soil to the cabinet air outside the shoot chamber through a single Keeling plot, i.e., by plotting cabinet ^13^C-CH_4_ over its inverse concentration 24 h after injection. The proportion of CH4 emitted by shoots of S. lapponum originated from peat was calculated as:

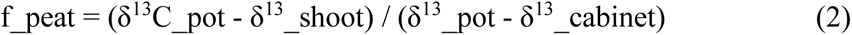

Where f_peat is the proportion of ^13^C-CH4 emitted by shoots from peat, δ^13^C_pot, δ^13^_shoot and δ^13^_cabinet are the value of δ^13^ in pot, shoots, and cabinet, respectively.

### Targeted metagenomic analysis of CH_4_ cycling microbes

To evaluate the possibility of microbial processes within or on the surfaces of the plants contributing to the plant-derived CH_4_ flux, we conducted a probe-targeted metagenomic analysis (Siljanen *et al*., 2022) of *C. rostrata* (whole shoot), *M. trifoliata* (whole shoot, without flowers), *S. lapponum* (stem and leaves separately) and *B. nana* (shoot, i.e. branches with the leaves, and stem separately). The analysis targeted functional genes *mcrA*, coding for the methyl coenzyme-M reductase (methanogens); *pmoA*, coding for the particulate methane monooxygenase (methanotrophs); and *mmoX*, coding for the soluble methane monooxygenase (methanotrophs). Plant samples (n = 3 per sample type) were collected aseptically during the “high summer” campaign in 2021, except for *B. nana*, which was sampled already in August 2019 (n = 1). All samples were transported on ice and stored in -20 ^°^C until DNA extraction using the Nucleospin Plant II mini kit (Macherey-Nagel, Düren, Germany). Extraction included mortar homogenization in liquid nitrogen and was conducted according to the manufacturer instructions, except of the extension of the lysis step to 1 hour. DNA quantity and quality was checked with Qubit fluorometer and Nanodrop One spectrophotometer (both Thermo Fisher Scientific). Targeted sequencing was conducted by Daicel Arbor Biosciences (US) as in Putkinen *et al*. (2021). Data was analysed as in Putkinen *et al*., 2021, except for the jplace-file processing and visualization, which was done with Gappa (Czech *et al*., 2020). Reagent and sample cross-contamination was controlled via parallel sequencing of blank-samples (only DNA-extraction reagents, no plant material). The sequence data produced in this study was deposited to the SRA-NCBI database under the BioProject PRJNA953003.

### Statistical analyses

To investigate the effects of seasonality, temperature and clipping treatment on the flux under standardized [CH_4_]_pw_ conditions, fCH_4p_ was normalized by [CH_4_]_pw_. This produced a variable hereafter called “apparent plant CH_4_ transport efficiency” (fCH_4pe_) as we cannot exclude that relevant amounts CH_4_ were produced or oxidized within the plants. When evaluating the effect of PAR, we assumed that diurnal changes in [CH_4_]_pw_ (measured only once per day) and its impacts on fCH_4p_ were negligible. Thus, we did not investigate the effect of PAR on fCH_4pe_.

The effect of phenology on fCH_4p_ and fCH_4pe_, as indicated by the differences among mesocosms collected at the three different plant phenological phases over the two growing seasons, was tested with one-way ANOVA and post hoc tests. We used linear mixed-effects models (LMMs) to examine the effects of PAR and [CH4]pw on the fCH4p, as well as the effects of temperature on the fCH4p and fCH4pe. The LMMs was also used to test the effects of clipping treatment on the fCH4p and fCH4pe of C. rostrata. These tests were conducted for each species separately. Fixed predictors, analysed one by one, each in a separate LMM, were the environmental variables [CH_4_]_pw_, temperature (15 vs 20 °C) and PAR (2 levels of PAR) as well as clipping treatment (intact vs shoot clipping), and the random variable was sample id in all LMMs analyses. The models are presented in Table 2, S2-S4. No statistics was applied for the number of the genes of the microbiology data. All statistic analyses were performed using R v.3.6.1 (Team, 2019).

**Table 2.** Summary statistics for the linear mixed-effects model describing the effects of porewater CH_4_ concentration ([CH_4_]_pw_) on the plant CH_4_ flux (fCH4p) of *C. rostrata*, *M. trifoliata*, *B. nana* and *S. lapponum*. The observations used here are before the clipping and labelling experiments. Asterisks denote statistical significance: **, 0.01; ***, 0.001.

| *C. rostrata* |  |  |  | *M. trifoliata* |  |  |  |
| --- | --- | --- | --- | --- | --- | --- | --- |
| Fixed part | Estimates | SE | P value | Fixed part | Estimates | SE | P value |
| Intercept | 1.45889 | 1.95450 | 0.48500 | Intercept | 0.11557 | 0.11557 | 0.29800 |
| [CH<sub>4</sub>]<sub>pw</sub> | 0.03266 | 0.00422 | < 0.001 *** | [CH<sub>4</sub>]<sub>pw</sub> | -0.00085 | 0.00018 | < 0.001 *** |
| Random part |  |  |  | Random part |  |  |  |
| SD (Sample ID) | 19.13100 | 4.37400 |  | SD (Sample ID) | 0.03371 | 0.18361 |  |
| Residual SD | 3.52400 | 1.87700 |  | Residual SD | 0.00355 | 0.05957 |  |
| *S. lapponum* |  |  |  | *B. nana* |  |  |  |
| Fixed part | Estimates | SE | P value | Fixed part | Estimates | SE | P value |
| Intercept | -0.00307 | 0.00146 | 0.12385 | Intercept | 0.13380 | 0.09446 | 0.21800 |
| [CH<sub>4</sub>]<sub>pw</sub> | 0.00003 | 0.00001 | < 0.01 ** | [CH<sub>4</sub>]<sub>pw</sub> | 0.00009 | 0.00033 | 0.79500 |
| Random part |  |  |  | Random part |  |  |  |
| SD (Sample ID) | 0.00001 | 0.00289 |  | SD (Sample ID) | 0.00015 | 0.03906 |  |
| Residual SD | 1.808e-06 | 0.00148 |  | Residual SD | 3.801e-06 | 0.00094 |  |

## Results

### Seasonality of porewater concentration and plant CH_4_ flux

The CH_4_ concentration ([CH_4_]_pw_) of C. rostrata and B. nana mesocosms showed wide variability from the minimum to the maximum (Fig. S1). The minimum was 0.02 µmol l^-1^ for C. rostrata and B. nana mesocosms, both were over 10000 times lower than that for other growing seasons. Contrastingly, the variability from the minimum to the maximum [CH_4_]_pw_ in *M. trifoliata* and *S. lapponum* mesocosms was small (Fig. S1). The [CH_4_]_pw_ was lowest high summer, while highest in early autumn, with means of 17 and 40 µmol l^-1^ for M. trifoliata mesocosms, and 21 and 23 µmol l^-1^ for S. lapponum mesocosms, respectively. [CH_4_]_pw_ significantly increased the fCH4p of *C. rostrata* and *S. lapponum* and decreased the fCH4p of *M. trifoliata*, (all P < 0.01, Table 2). In contrast, no significant relationship between porewater concentration and plant flux was observed for *B. nana*.

To focus on the seasonality unrelated to [CH_4_]_pw_ we investigated the variability in the apparent transport efficiency (fCH_4pe_). The seasonal pattern in the apparent transport efficiency (fCH_4pe_) and in the plant CH_4_ flux (fCH4p) appeared to differ between the four studied species. The fCH_4pe_ of *C. rostrata* was higher (mean 234 mg CH_4_ m^-2^ leaf area h^-1^ (µmol l^-l^)^-1^ in high summer than in early summer and autumn (0.02 and 0.06 mg CH_4_ m^-2^ leaf area h^-1^ (µmol l^-l^)^-1^, respectively) (both P < 0.001, Fig. 2). However, the mean fCH_4p_ of *C. rostrata* was at its minimum in early summer (3.04 mg m^-2^ h^-1^), increased to 4.86 mg m^-2^ h^-1^ in high summer, and reached its maximum in early autumn (10.70 mg m^-2^ h^-1^). Similar to *C. rostrata*, the maximum transport efficiency of *B. nana* occurred in high summer (mean 4.08 mg m^-2^ h^-1^ (µmol l^-l^)^-1^) but the difference to the early summer minimum (0.001 mg m^-2^ h^-1^ (µmol l^-l^)^-1^) for *B. nana* was smaller in magnitude than in *C. rostrata*. The maximum mean plant flux of *B. nana* occurred in early summer (0.73 mg m^-2^ h^-1^) and the minimum in early autumn (0.001 mg m^-2^ h^-1^), respectively. For *M. trifoliata*, both transport efficiency and plant flux were significantly higher in high summer than in early autumn (both P < 0.001). For *S. lapponum*, both transport efficiency and plant flux were significantly smaller in early and high summer than in early autumn (both P < 0.001).

**Fig. 2.**
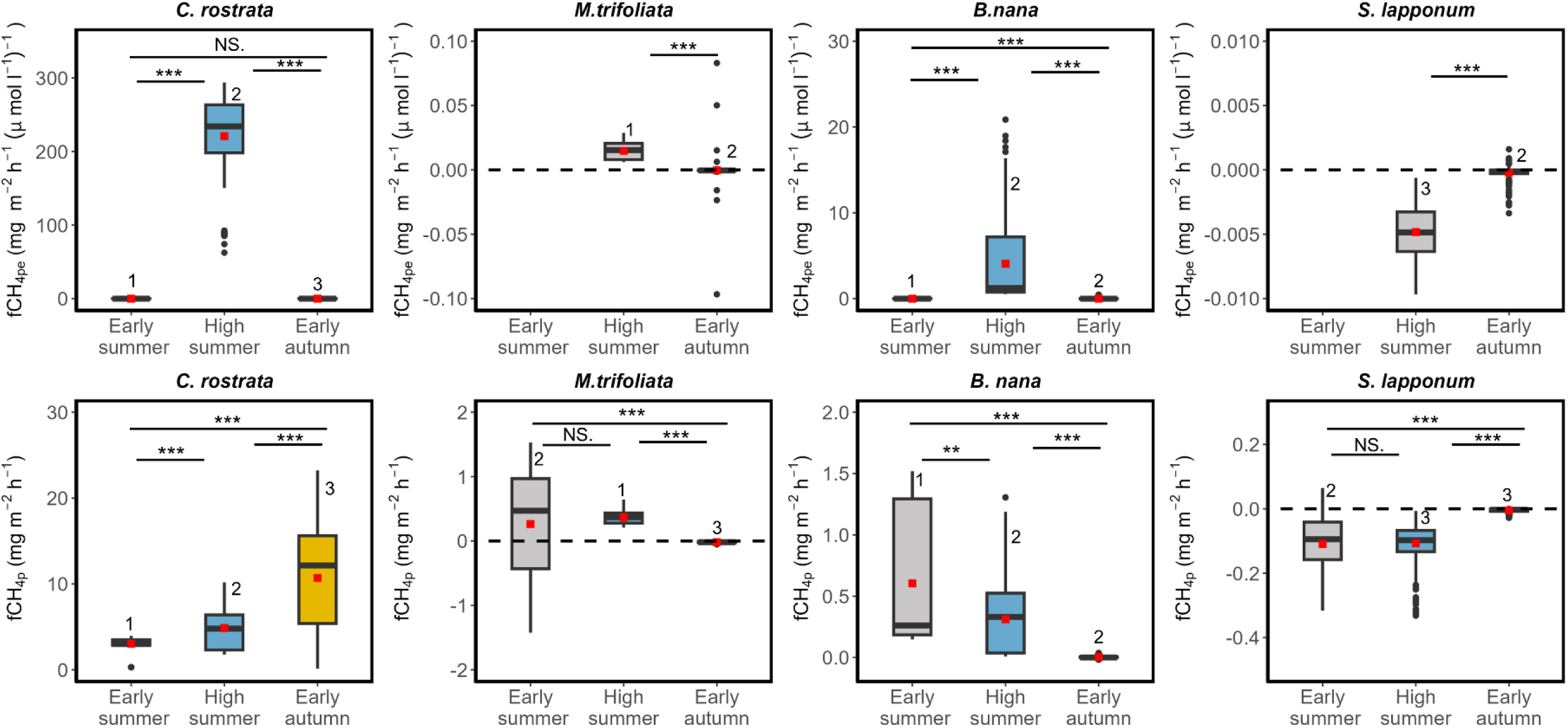
Characteristics of the seasonal apparent CH_4_ transport efficiency (fCH_4pe_, mg CH_4_ m^-2^ leaf area h^-1^ (µmol CH4 l^-l^ porewater)^-1^)) and plant flux (fCH_4p_, mg CH_4_ m^-2^ leaf area h^-1^) for herbs *C. rostrata* and *M. trifoliata*, and shrubs *B. nana* and *S. lapponum*. Red dots are mean fCH_4pe_ and fCH_4p_ of each season. Note the different y-axis scales. The number above the box denotes the number of specimens observed during each campaign. NS denotes no significant difference and asterisks denote statistical significance: **, 0.01; ***, 0.001.

### Effects temperature and PAR

The response to increase in temperature from 15°C to 20°C, differed between the plant species and also between apparent CH_4_ transport efficiency and plant methane flux (Fig. S4). The transport efficiency of *C. rostrata* significantly decreased after the temperature was increased (P < 0.001, Fig. 3) while in the other three species there was no response. However, with the plant flux calculated per gram of dry mass, higher temperature leaded to higher flux of *C. rostrata* and *S. lapponum* (both P < 0.001), while it reduced the flux of *M. trifoliata* (P < 0.001). No significant temperature effect was observed in the flux of *B. nana*. As for the effects of PAR, it did not significantly affect fCH_4p_ of any of the investigated species (Fig. 4).

**Fig. 3.**
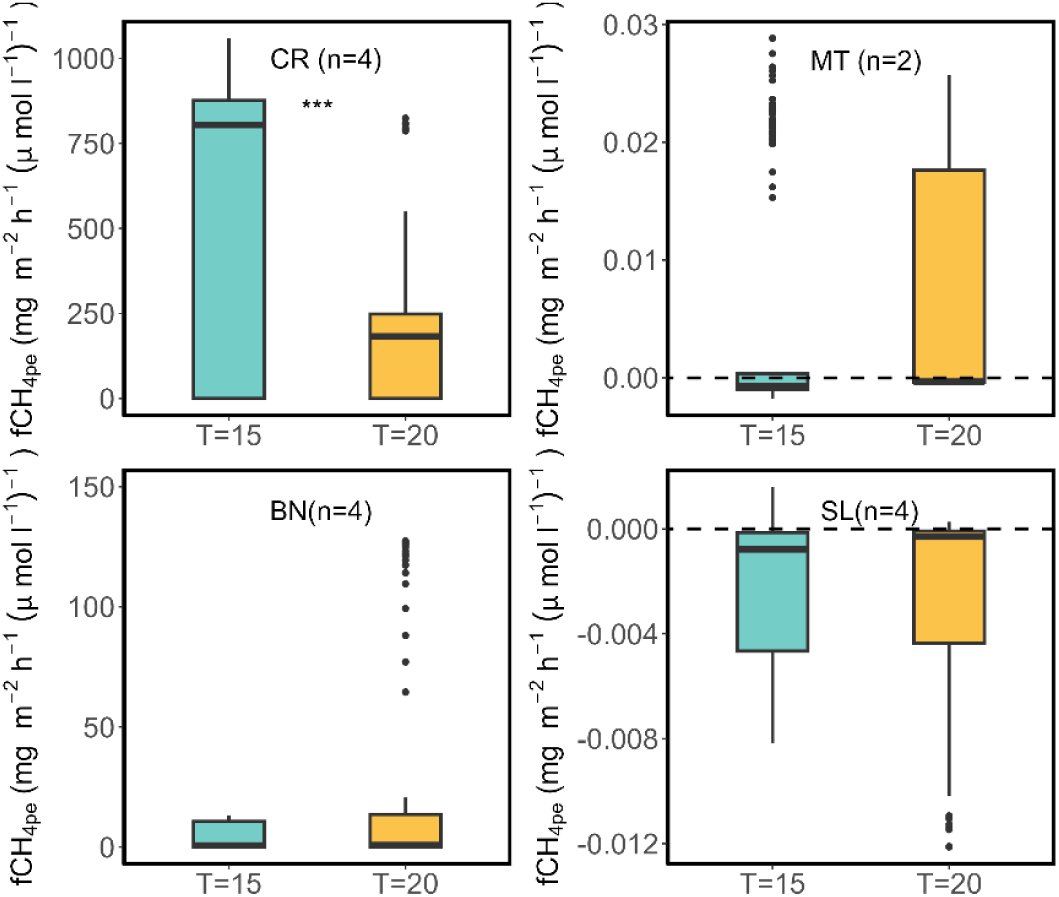
Temperature effects on apparent CH_4_ transport efficiency (fCH_4pe_, mg CH_4_ m^-2^ leaf area h^-1^ (µmol l^-l^)^-1^ of *C. rostrata* (CR), shrubs *B. nana* (BN) and *S. lapponum* (SL). Summary statistics for the linear mixed-effects model describing the effects of temperature on fCH_4pe_ shown in Table S2. The number (n) of total mesocosms per species is less than 9 because of system and GC malfunctions. Dashed horizontal line in panels with negative fCH_4pe_ denotes zero flux level. Note the different y-axis scales. Asterisks denote statistical significance: ***, 0.001.

**Fig. 4.**
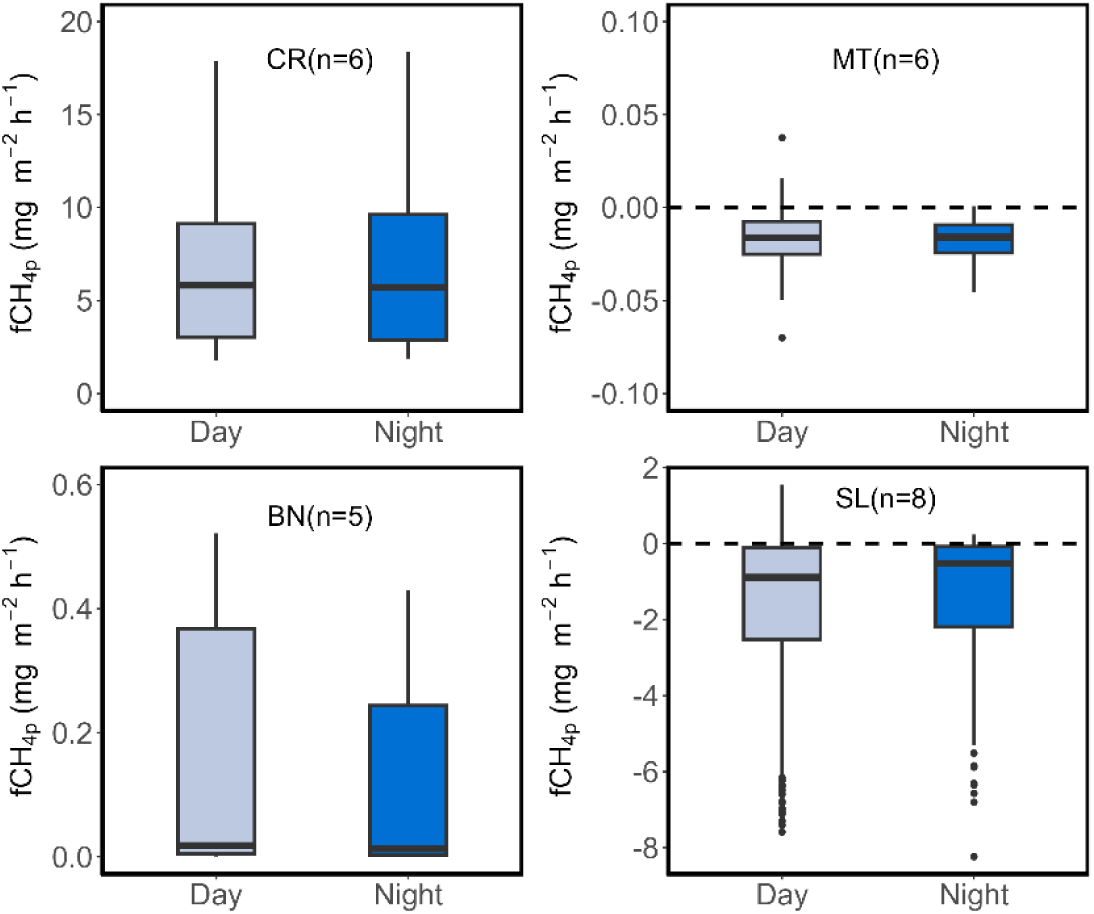
PAR effects on plant CH_4_ flux (fCH_4p_, mg CH_4_ m^-2^ leaf area h^-1^) of *C. rostrata* (CR)*, M. trifoliata* (MT), and *B. nana* (BN) and *S. lapponum* (SL). The number (n) of total mesocosms per species is less than 9 because of system and GC malfunctions. Dashed horizontal line in panels with negative fCH_4p_ denotes zero flux level. Note the different y-axis scales.

**Fig. 5.**
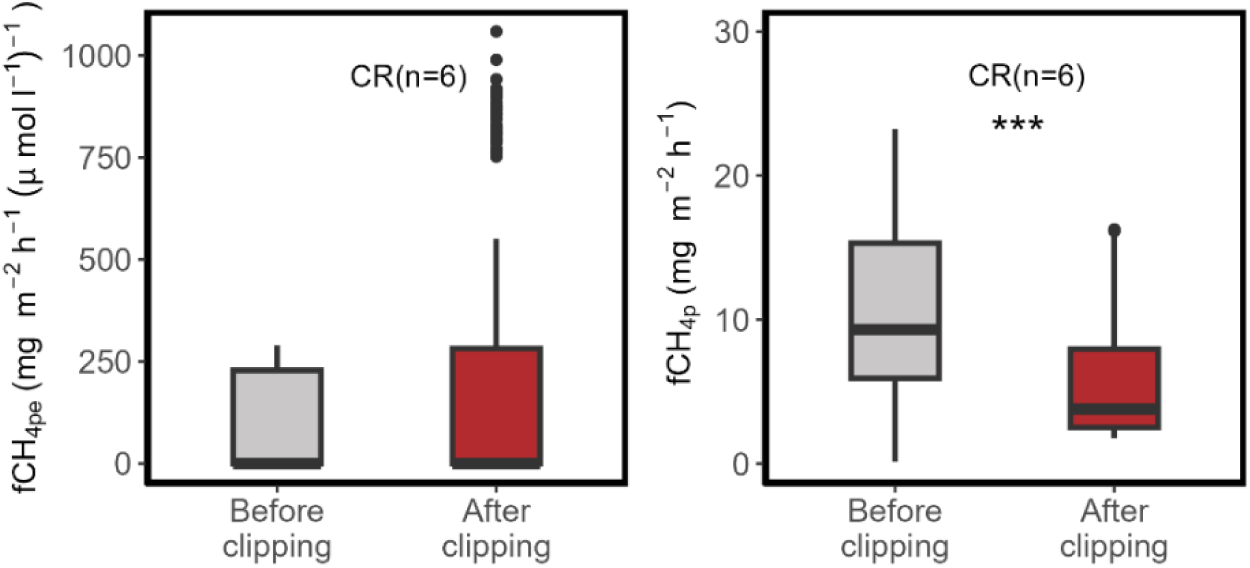
Clipping effects on apparent CH_4_ transport efficiency (fCH_4pe_, mg CH_4_ m^-2^ leaf area h^-1^ (µmol l^-l^)^-1^) and CH_4_ flux (fCH_4p_, mg CH_4_ m^-2^ leaf area h^-1^) of *C. rostrata* (CR). The number (n) of total mesocosms per species is less than 9 because of system and GC malfunctions. Same mesocosms before and after the clipping treatment. Asterisks denote statistical significance: ***, 0.001.

### Plant-associated methanogens and methanotrophs

We found some of the plant shoots to host microbes that produce or oxidize CH4 but not from all species. Targeted metagenomic sequencing, and the following phylogenetic placement analysis, revealed methanogenic and methanotrophic functional genes in the shoots of *M. trifoliata* and in the leaves and stem of *S. lapponum* (Fig. 8, Figs. S6‒S8). None of the genes were detected in *C. rostrata*. In the separately screened *B. nana* samples (shoot and stem, n =1), methanotrophs, but no methanogens, were found (Fig. 9). Sequence read numbers (after the hmmer profile screening) were low with mean counts in different sample types varying from 0 to 378 reads in *mcrA*, from 48 to 393 in *pmoA* (including *amoA*), and from 7 to 10782 in *mmoX* (including non-methane mono-oxygenases; Table S3).

**Fig. 7.**
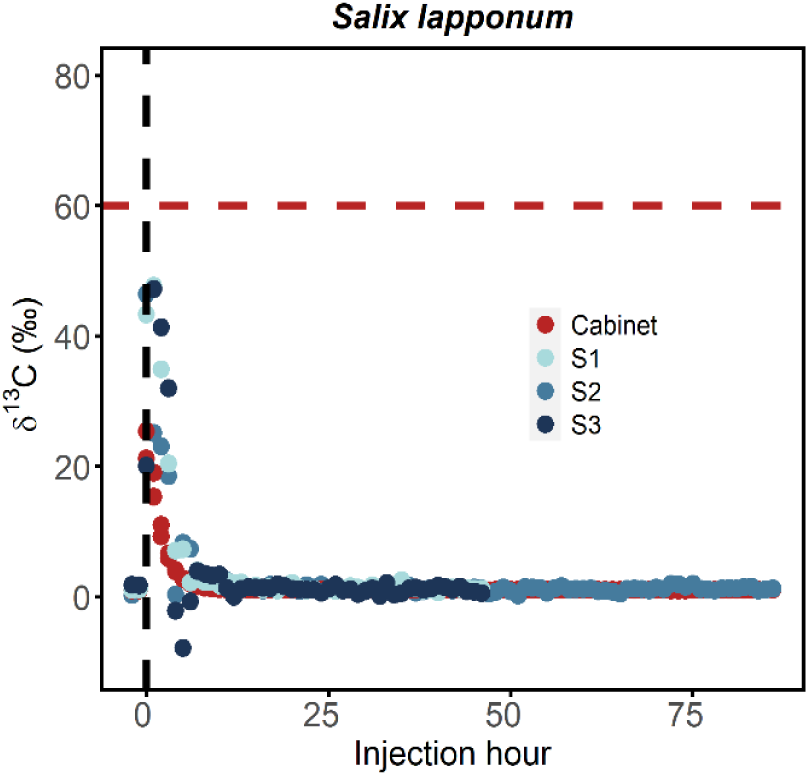
^13^C-CH_4_ labelling result of CH_4_ fluxes through *S. lapponum*. Red line is the intercept of keeling plot, representing δ^13^C-CH_4_ *S. lapponum* mesocosms (S1, S2 and S3). Dashed vertical black line represents the moment of injection. Red dots are the δ^13^C-CH_4_ of the cabinet where mesocosms were placed. Blue dots with different shades are the δ^13^C-CH_4_ of fluxes emitted by shoots of *S. lapponum*.

**Fig. 8.**
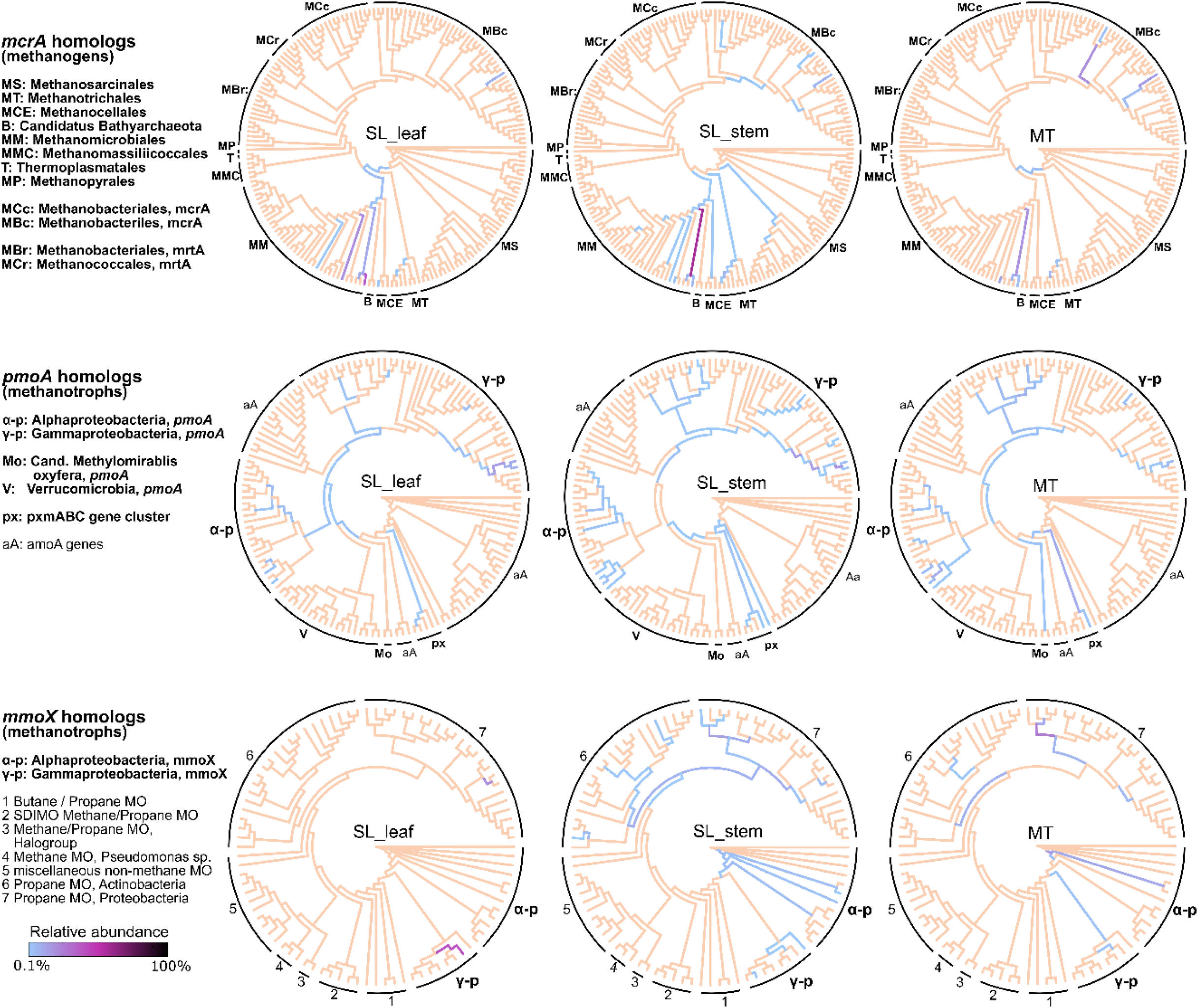
Diversity of CH_4_ cycling genes *mcrA* (for methanogens), *pmoA* (for methanotrophs) and *mmoX* (for methanotrophs) in *S. lapponum* (SL) leaf and stem samples, and in *M. trifoliata* (MT), combined stem and leaf (n =3). Taxonomy of the gene sequences was analysed via phylogenetic placement on the reference trees of genes from known strains (placements accumulated to the inner branches based on the likelihood-weight threshold of 0.8). Figure shows these placements as heat-trees, where relative abundances of placements are visualized as indicated by the colour bar. Reference branches with no placements or <0.1% relative abundance are shown with light orange. Composition of the reference strains reflects the probes included in our analysis = coverage of the analysis. Detailed information of each of the reference strains can be found in Figs. S7-S9. Results for *B. nana* are in Fig. 8. No sequences were detected in *C. rostrata*.

**Fig. 9.**
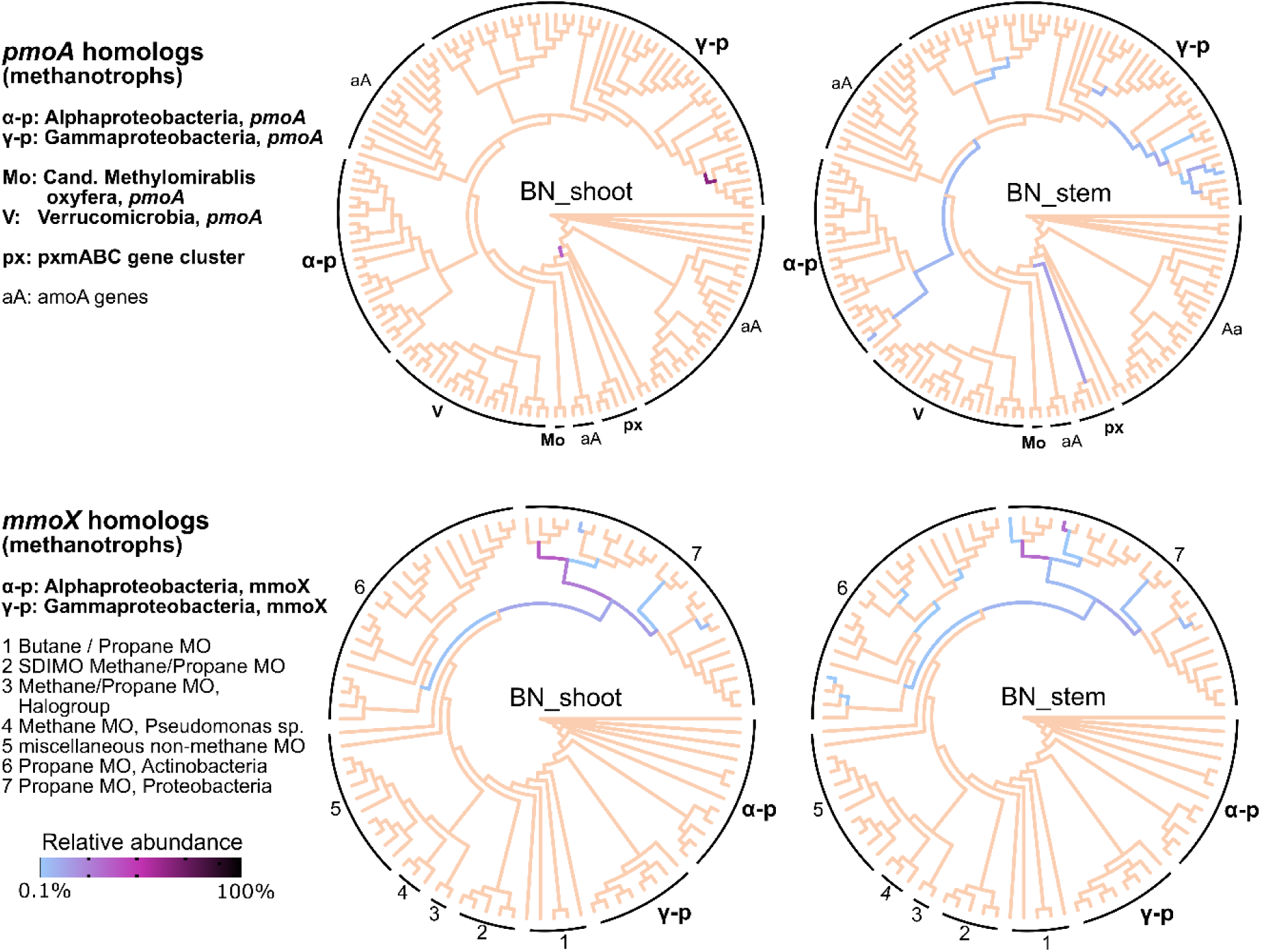
Diversity of methanotrophic genes *pmoA* and *mmoX* in *B. nana* (BN) shoot and stem samples (n =1). Taxonomy of the gene sequences was analysed via phylogenetic placement on the reference trees of genes from known strains (placements accumulated to the inner branches based on the likelihood-weight threshold of 0.8). Figure shows these placements as heat-trees, where relative abundances of placements are visualized as indicated by the colour bar. Reference branches with no placements or <0.1% relative abundance are shown with light orange. Composition of the reference strains reflects the probes included in our analysis = coverage of the analysis. Detailed information of each of the reference strains can be found in Figs S8 and S9. No *mcrA* sequences were detected in *B. nana*.

In both *M. trifoliata* and *S. lapponum,* sequences of the *mcrA* gene were mainly related to the genera *Methanobacterium (order Methanobacteriales)* and *Methanoregula (Methanomicrobiales)*, accompanied by *Methanotrichales* and *Methanocellales*. The highest diversity of different genera was detected in the stems of *S. lapponum*.

In *M. trifoliata,* methanotrophic *pmoA* genes were related to both alpha- and gammaproteobacteria, with *Methylocystis* & *Methylocapsa* and *Methylobacter* & *Methylomagnum* as the most abundant genera within those classes, respectively. In *S. lapponum, Methylocystis* and *Methylobacter* were the largest groups. *B. nana* harboured alpha- (in shoot & stem) and gammaproteobacterial (in stem) *pmoA* genes. All sample types, except *C. rostrata*, showed presence of the still poorly known *pxmABC* gene cluster (Tavormina *et al*., 2011) and also of alphaproteobacterial Upland soil cluster α, common in habitats with low/atmospheric CH_4_ concentrations (i.e. capable of high-affinity oxidization) (Tveit *et al*., 2019; Fig. S6). Based on the *mmoX* gene (found only in some of the methanotrophic taxa) *M. trifoliata* and *S. lapponum* stem contained both alpha- and gammaproteobacterial methanotrophs (especially *Methylocella* and family *Methylococcaceae*). *S. lapponum* leaf contained only gammaproteobacterial *mmoX* (*Methylomagnum*), and no *mmoX* was present in the *B. nana* samples, only genes for other related mono-oxygenases.

### Impact of Carex rostrata clipping

Clipping treatment where leaves were removed but the stem was left untouched did not significantly affected the apparent CH_4_ transport efficiency of *C. rostrata* (Fig. S3). However, higher plant CH_4_ flux was observed.

### Source of plant CH_4_ flux from Salix lapponum

The isotope labelling revealed that methane injected into the root space is rapidly transported through *S. lapponum* and emitted from its shoot (Fig 7). After the injection, the δ^13^C-CH_4_ value of mesocosm CH4 (inferred from the CH4 emitted into the cabinet air) was *c.* +60‰ (red line in Fig. 7). Assuming a background δ^13^C-CH_4_ value of -50‰, our injection has therefore changed [CH4]_pw_ less than 0.15% of the concentration prior labelling, i.e., the label application had minimal effects of the environment of S. lapponum roots. During the measurements, the maximum δ^13^C-CH_4_ in the cabinet was *c.* +26‰, which was lower than that from the mesocosm due to mixing with ambient air. δ^13^C-CH_4_ emitted into shrub chambers was *c.* +48‰, which was higher than that of the cabinet air (*c.* +26‰) and closer to δ^13^C-CH_4_ in the mesocosm (*c.* +60‰). Labelling also showed that major part of CH_4_ (*c.* two thirds) emitted from *S. lapponum* was transported from the mesocosm through plant and emitted through the shoot, rather than from leakage of cabinet air into the shrub chamber.

## Discussion

We quantified the magnitude and seasonality of the CH_4_ flux from shoots of four common boreal peatland plants, and the responses of the flux to temperature, PAR and porewater CH_4_ concentration ([CH_4_]_pw_) using a high-frequency, automated, climate-controlled measurement system, The species represented different plant functional types, including a sedge (*Carex rostrata*), a forb (*Menyanthes trifoliata*), and two shrubs (*Salix lapponum* and *Betula nana*). Further, we used the clipping treatment to reveal the role of leaves in restricting the emission from *C. rostrata,* and labelling and microbial analyses to evaluate the source of the plant-emitted CH_4_. This is to our knowledge the first study to disentangle the importance of phenology and abiotic factors in regulating CH_4_ fluxes through peatland plants and to examine the presence of CH_4_ cycling microbes in a range of peatland plants. We found that phenology is an overriding factor in controlling plant-mediated CH_4_ flux, but that the magnitude and seasonal course of phenology effects is species-specific. Out of the studied abiotic factors, temperature and porewater CH4 concentration were also found to regulate plant-mediated CH4 flux, but the magnitude and direction of the effect of these variables differed between species. Hence, besides phenology, our results highlight the substantial role of species-specific differences in regulating the plant-mediated CH4 flux. Based on the results of our clipping experiment with C. rostrata, CH4 release was not regulated by leaves. The plant transported CH4 in the tested species, Salix lapponum, appeared to be mainly soil-derived. However, the discovery of methanogens and methanotrophs in the plant shoots suggests that the CH4 emission through some plants may also be modulated by CH_4_ production and oxidation by microorganisms inside the plant.

### Plant phenology controls plant-mediated CH_4_ flux

Our measurements were conducted in a climate-controlled environment, and therefore the strong seasonality observed in plant CH_4_ flux (fCH_4p_) was ascribed to phenology rather than seasonally different weather conditions. The fCH4p of C. rostrata, a widely studied species and the most efficient CH4 emitter, was significantly higher in high summer than that in early summer (Fig. 2), matching our earlier field measurement results (Ge et al., 2023). With this species, the variations in the transport between seasonal growth stages may be linked to the known seasonal changes in plant morphology. In early summer, newly-merged shoots do not have roots (Hultgren, 1989), and the aerenchyma size is rather small (Fagerstedt, 1992), which could explain the small fCH4p in early summer. In contrast, fCH4p of M. trifoliata and S. lapponum did not change between early and high summer, which suggests that the aboveground leaf-out during this phase would not affect their CH4 transport. With B. nana, fCH4p was higher in early summer than that in high summer.

The smaller amount of shoot biomass in early summer than that in high summer might also cause the higher fCH4p, yet the possibility is small due to following reasons. Firstly, no correlation between leaf area and CH4 flux from C. rostrata was observed in this study (Fig. S9) or the field measurement (Ge *et al*., 2023). Secondly, clipping the shoots did not significantly change the fCH4pe of C. rostrata (Fig. 6). Thirdly, stomatal control in aerenchymatous plants is poorly developed due to living in high-moisture environments (Lange *et al*., 1971). This echoes with the poor correlation between PAR and the fCH4p of C. rostrata and other investigated species and also the fact that CH4 can be released from stems or leaf sheaths, as found in many species like rice plants (Nouchi *et al*., 1990) and forbs (Shannon *et al*., 1996). All these lead us to believe that seasonal changes in belowground parts of C. rostrata, instead of leaves, control CH4 transport of C. rostrata, which is against our hypothesis. Unlike C. rostrata, the fCH4p of B. nana was higher in early summer than that in high summer. However, due to the lack of morphological data, it is unknown what happens for them in this par of the growing season leading the changes in the fCH4p.

The fCH4p of C. rostrata was higher in early autumn when it was senescencing than that in high summer. This indicates the increasing proportion of brown leaves does not affect the transport of C. rostrata, which is again support the speculation that the belowground parts of plant regulate the transport. The root permeability, a key parameter controlling plant CH4 transport capacity (Beckett et al., 2001; Henneberg et al., 2012), could decrease in early autumn (Nouchi *et al*., 1994; Wassmann & Aulakh, 2000) and, thus, reduce the fCH4p of C. rostrata. Yet, the significantly higher porewater CH4 concentration in early autumn than that in high summer might compensate for the effect of root permeability on the transport and lead to the maximum fCH4p.

In early autumn, M. trifoliata was senescencing and shrubs B. nana and S. lapponum had already dropped all their leaves. For M. trifoliata and B. nana also the CH4 flux decreased from high summer to early autumn. As for S. lapponum who constantly showed an uptake of CH4, and the remarkable increase of fCH4p in early autumn means the decrease of absorption rate. The changes in the fCH4p of these species in early autumn might indicate that physiological activity, or lack of leaves (for shrubs only), might be the key behind the phenology-driven CH4 flux in these species. The importance of physiology and leaves in regulating CH4 transport of herbs and woody species has also been reported (Garnet et al., 2005; Pangala et al., 2014; Pangala et al., 2015). Yet, CH4 flux via these species did not respond to PAR, and the effects of foliage removal to the flux are still unknown. Also, the seasonal changes in the belowground parts of these species are unknown, making us unable to identify the main control behind phenology in regulating CH4 flux.

### Porewater CH_4_ concentration affects CH4 flux from most investigated species

We found that [CH_4_]_pw_ increased the fCH_4p_ of *C. rostrata* and S. lapponum, decreased the flux of *M. trifoliata*, but had no effect on *B. nana* (Table 2), even though [CH_4_]_pw_ varied in all mesocosms (Fig. S1, S2). As an indicator of CH_4_ supply to the roots, [CH_4_]_pw_ has been reported to control CH_4_ flux from both herbs (Schimel, 1995; Aulakh *et al*., 2000), and woody species (Pangala *et al*., 2014). Yet, in our earlier *in situ* study we did not observe any relationships between the fCH_4p_ and [CH_4_]_pw_ for any of the studied species (Ge *et al*., 2023). Howeverm, in this field study, [CH_4_]_pw_ varied less (from 3.33 to 537 µmol l^-1^, Ge *et al*., 2023) than the concentrations in the current laboratory study (from 0.0018 to 511 µmol l^-1^),. Notably, CH_4_ flux from *C. rostrata* did not show any signs of saturation even though [CH_4_]_pw_ (Fig. S10) was high in early summer and autumn (mean 105 and 284 µmol l^-1^, respectively). In contrast, CH_4_ flux through rice reached a saturation point when [CH_4_]_pw_ reached merely 14 µmol l^-1^ (Aulakh *et al*., 2000). This indicates that the ability of plants to increase their transport with higher [CH_4_]_pw_ is species-specific. Species reaching the saturation point faster could then potentially limit the total CH4 flux into the atmosphere.

The close relationship between [CH_4_]_pw_ and the fCH_4p_ of *M. trifoliata* (Table 2) suggest that [CH4]pw controls the fCH4p of *M. trifoliata*. This result implies that the high fCH4p of M. trifoliata observed in the field (Ge et al. 2023) was probably caused by the constantly and significantly high [CH4]pw where it grew. In contrast, the fCH4p of M. trifoliata was small in the climate-controlled measurement in the present study where [CH_4_]_pw_ in *M. trifoliata* mesocosms were small, up to 27 times lower than that in the field. Thus, instead of a high transport efficiency, the high fCH4p of M. trifoliata observed in the field could be mainly due to the high [CH_4_]_pw_.

### Temperature decrease plant CH4 flux, but only from C. rostrata

The fCH_4pe_ of *C. rostrata* significantly decreased after increasing temperature, reflecting that a higher temperature stimulated both porewater CH4 production (increasing the [CH_4_]_pw_) and fCH_4p_, but the increase of [CH_4_]_pw_ was relatively higher than the increase of fCH_4p_, leading to a lower CH4 transport efficiency (fCH_4pe_). Thus, potentially, the ability of *C. rostrata* to transport CH_4_ could be a limiting factor for ecosystem CH_4_ emissions when rising temperatures and higher substrate availability stimulates methanogenesis. As the fCH4pe of the other studied species did not respond to warmer temperature, it appears that the temperature effect on plant transport is species-specific. The negative impacts of peat temperature on the transport of *C. rostrata* were also observed in our situ measurement conducted in the high summer when peat temperature was at a similar range as in this climate cabinet study (11 °C to 17 °C) (Ge *et al*., 2023).

It is evident that an experiment in controlled climate cabinet cannot, and by definition should not, directly mimic natural conditions and hence the interpretation of the temperature dependency should be considered with care. The decreasing transport of C. rostrata after the increase of temperature could also be due to decreased activity or increased stress of the plants inside the cabinet. However, we did not observe a significant decrease of net ecosystem exchange, an indicator of plant growth conditions, after the increase of temperature (Fig. S10), which suggests that the plants would not have experienced at least a severe stress. We also acknowledge that the field sampling, transport and setting up of the experiment can have caused disturbance to the plants. However, the comparable fCH4p measured in the field (-5 to 25 mg CH_4_ m^-2^ leaf area h^-1^) and in the climate-controlled cabinet (-0.4 to 21 mg m^-2^ h^-1^) suggest that the plants displayed rather realistic transport rates in the laboratory conditions. Further, the temperature and light conditions were chosen to represent typical field conditions to which the plants have adapted to.

### CH_4_ exchange processes of plants include within-plant CH4 production and oxidation

Role of plant-associated microbes in peatland CH_4_ cycle is still poorly understood, apart from the association between methanotrophs and *Sphagnum* mosses (e.g. Larmola *et al*., 2010; Putkinen *et al*., 2014). Especially, to our knowledge, markers of microbial CH_4_ production within plant tissues has never been reported in this context. Based on our results, methanogenic archaea can be a part of the microbiome of both forbs (*M. trifoliata)* and shrubs (*S. lapponum*, in both stem and leaf parts), which suggests that microbial CH_4_ production could occur in the shoots of these plants.

The detected methanogens included both hydrogenotrophic (*Methanocellales*, *Methanomicrobiales, Methanobacteriales)* and acetoclastic (*Methanotrichales*) orders (Knief, 2019). Interestingly, most of them (*Methanocellales*, *Methanomicrobiales* and *Methanotrichales*) are proposed to possess features allowing adaptation to oxidative environments (Lyu & Lu, 2018) – such as above-ground plant parts. Our findings corroborate previous plant microbiome studies: *Methanomicrobiales* and *Methanotrichales* were found in spruce needles (Putkinen *et al*., 2021) and *Methanomicrobiales, Methanocellales and Methanobacteriales* as part of poplar stem wood (Yip *et al*., 2019; Feng *et al*., 2022). Yet, all these taxa are commonly found also in the anaerobic peat, including Lompolojänkkä where our plant-soil mesocosms were collected (unpublished, Putkinen *et al*).

Similar to methanogens, methanotrophs were discovered in forb (*M. trifoliata*) and in both studied shrubs (*S. lapponum*, *B. nana*). This result implies potential for CH_4_ oxidation within these plants that have long been merely regarded only as CH4 transporters (Shannon *et al*., 1996; Ding *et al*., 2005; Ge *et al*., 2023). The detection of methanotrophs inside shrub S. lapponum fits well with the plant as a whole acting as a constant CH4 sink (negative fCH_4p_) throughout the growing seasons (Fig. 2). Although B. nana mostly emitted CH_4_ in this study, our earlier field measurements of *B. nana* showed consumption of CH_4_ in early summer (Ge *et al*., 2023), and Riutta *et al*. (2020) also reported the attenuating effects of B. nana on CH4 flux.

The methanotrophs we detected entailed several taxonomic groups, representing both obligate and facultative methanotrophs (latter found mainly in the alphaproteobacteria (Knief, 2015)). Diversity was demonstrated also in the presence of variable forms of methane mono-oxygenases: in addition to common particulate (pMMO, coded by *pmoA*) and soluble (sMMO, coded by *mmoX*) forms, all plants, except *Carex*, contained genes for the pxmABC cluster, thought to code a novel type of particulate MMO, the pXMO (Tavormina *et al*., 2011). Although the function of the pXMO is still poorly understood, it has been suggested to support methanotrophs under hypoxia (Kits *et al*., 2015). On the species-level, *Methylocystis bryophila*, able to use multiple C sources and entailing pMMO variants for both low-(pMMO1) and high-affinity (pMMO2) CH_4_ oxidation (Han *et al*., 2018), highlights the versatility of the detected methanotrophs. In addition, sequences of the USCa, including of the first cultivated organism from this group, *Methylocapsa gorgona* (Tveit *et al*., 2019), further demonstrated that plant-associated methanotrophs may affect peatland CH_4_ balance not only by consuming soil-derived CH_4_ but also as a sink for the atmospheric CH_4_. Especially *Methylocystis* methanotrophs have been detected in other plant types as well (Putkinen *et al*., 2021), and like with methanogens, most methanotrophs living in Lompolojänkkäplants have close relatives in the surrounding peat (unpublished, Putkinen *et al*.) and in other boreal peatlands.

Our results indicate that within-plant microbial control mechanisms could play a role in modulating plant-derived CH_4_ flux, particularly by the consumption of CH_4_ transported from the soil inside the plant, indicated by the negative fCH4p of S. lapponum. However, quantification of these microbes and the related processes is challenging. This stems from the methodological difficulties in extracting the endo- and epiphytic microbes from among the vast amounts of plant-derived genetic material. Novel tools, like probe-targeting, do aid in this task, but still leave room for uncertainties (e.g., regarding sensitivity of the analysis in different plant material types). This, together with the generally low number of recovered sequences, limits the quantitative comparison of microbial communities in differing samples. Moreover, for the evaluation of their actual activity, analyses need to go beyond the DNA level and require additional analysis, e.g. visualisation of microbes within the plant structures or at least analysing samples where surface microbes would have been rinsed away).

## Conclusion

The number of high-frequency observations (around 4500 flux measurements in total), make our data unique and give a way forward to disentangling the roles of plant phenological stages and environmental drivers in plant-mediated CH4 fluxes. Our results show that plant CH4 emission and uptake varies between species and with plant growth stages over the growing seasons. Our study further shows that the responses of season-specific plant CH_4_ flux to temperature and porewater CH4 concentration vary between species. We detected methanogens and methanotrophs in the shoots of both herbs and shrubs, suggesting that in addition to transportation of soil-produced CH_4_, the plant-mediated CH4 flux could also involve within-plant CH_4_ oxidation and/or production. Further studies are needed to assess their roles and activities in different plant-soil environments, conditions and seasons.

## Author Contributions

AL, PM, MP, and MK designed the field study. MG conducted the field and laboratory measurements with methodological help from AP, LK, HS. MG, MK, AK, AP, HS analysed data. MG, MK, AK and AP wrote the manuscript with contributions from all other authors.

## Acknowledgements

We thank Rauna Lilja, Tatu Polvinen, Salla Tenhovirta for help with the climate-controlled measurements, Asko Simojoki for gas chromatograph instructions. The study was funded by the China Scholarship Council (CSC), the Academy of Finland (315415, 339489, 342362), the H2020 Marie Skłodowska-Curie Actions (843511), and H2020 European Research Council (757695).

## Data Availability

The original data are available at https://doi.org/10.5281/zenodo.7812992. Additional data related to this article can be obtained from the corresponding author (MG) on request.

