## Supplemental Table 1 for "CH_4_ transport in wetland plants under controlled environmental conditions – untangling the impacts of phenology"

New Phytologist

***New Phytologist* Supporting Information**

Article acceptance date: Click here to enter a date.

The following Supporting Information is available for this article:

**Table S1** Number of specimens for each species with available plant flux rate and porewater $CH_4$ concentration data.

**Table S2** Summary statistics for the linear mixed-effects model fitted to apparent $CH_4$ transport efficiency of *C. rostrata*, *B. nana* and *S. lapponum* with fixed effects of temperature.

**Table S3** Number of the sequence reads for each functional gene group.

**Figure S1** Porewater $CH_4$ concentration of *C. rostrata* and *B. nana* buckets measured in 2020.

**Figure S2** Porewater $CH_4$ concentration of *M. trifoliata* and *S. lapponum* buckets measured in 2021.

**Figure S3** Temperature effects on $CH_4$ flux of *C. rostrata*, *M. trifoliata*, shrubs *B. nana* and *S. lapponum*.

**Figure S6** Phylogenetic tree of 189 reference *mcrA* gene sequences used in the phylogenetic placement analysis of sequences retrieved from the plant samples.

**Figure S7** Phylogenetic tree of 142 reference *pmoA*/*amoA* gene sequences used in the phylogenetic placement analysis of sequences retrieved from the plant samples.

**Figure S8** Phylogenetic tree of 92 reference monooxygenase (MO) gene sequences used in the phylogenetic placement analysis of sequences retrieved from the plant samples.

**Fig. S9** Correlation between leaf area and $CH_4$ fluxes through *C. rostrata*.

**Fig. S10** Net ecosystem exchange of *C. rostrata* before and after the increase of temperature.

**Table S1** Number of specimens for each species with available plant flux rate and porewater $CH_4$ concentration data. Note 9 specimens in total of each species throughout the observations but end up with less than 9 for all species due to the system malfunctions (e.g., electricity problem, failure in controlling PAR and temperature or recording data), gas chromatograph malfunctions, and data quality control.

| | *C. rostrata* | *B. nana* | *M. trifoliata* | *S. lapponum* |
|---|---|---|---|---|
| Early summer | 1 | 1 | 0 | 0 |
| High summer | 2 | 2 | 1 | 3 |
| Early autumn | 3 | 2 | 2 | 2 |
| Total | 6 | 5 | 3 | 5 |

**Table S2** Summary statistics for the linear mixed-effects model fitted to apparent $CH_4$ transport efficiency of *C. rostrata*, *M. trifoliata*, *B. nana* and *S. lapponum* with fixed effects of temperature. Asterisks denote statistical significance: ***, 0.001.

| *C. rostrata* | | | | *M. trifoliata* | | | |
|---|---|---|---|---|---|---|---|
| **Fixed part** | Estimates | SE | P-value | **Fixed part** | Estimates | SE | P-value |
| intercept | 367 | 153 | 0.0953 | intercept | 0.0082675 | 0.009081 | 0.5298 |
| T=20 | -214 | 18 | < 0.001 *** | T=20 | -214 | 18 | 0.0703 |
| **Random part** | | | | **Random part** | | | |
| | Variance | SD | | | Variance | SD | |
| SD (Sample ID) | 92975 | 304.9 | | SD (Sample ID) | 0.00016486 | 0.01284 | |
| Residual SD | 18183 | 134.8 | | Residual SD | 0.00000222 | 0.00149 | |
| *B. nana* | | | | *S. lapponum* | | | |
| **Fixed part** | Estimates | SE | P-value | **Fixed part** | Estimates | SE | P-value |
| intercept | 25 | 22 | 0.323 | intercept | -0.00271389 | 0.001412 | 0.15 |
| T=20 | 0.2045 | 0.8678 | 0.814 | T=20 | 0.00008894 | 0.000127 | 0.484 |
| **Random part** | | | | **Random part** | | | |
| | Variance | SD | | | Variance | SD | |
| SD (Sample ID) | 2528 | 50 | | SD (Sample ID) | 7.941E-06 | 0.002818 | |
| Residual SD | 45 | 6.7 | | Residual SD | 2.214E-06 | 0.001488 | |

**Table S3** Number of the sequence reads for each functional gene group retrieved in the probe-targeted metagenome sequencing (n = 3, except for *B. nana*: n=1). Numbers are based on the target gene hmmer profile screening (E-value cut-off < 0.001).

| Gene | | Mean | Range |
|---|---|---|---|
| ***mcrA*** | *S. lapponum*, leaf | 41 | 0 – 112 |
| | *S. lapponum*, wood | 387 | 0 – 1156 |
| | *M. trifoliata* | 57 | 44 – 64 |
| | *B. nana*, shoot | 0 | - |
| | *B. nana*, stem | 0 | - |
| | | Mean | Range |
| ***pmoA***[1] | *S. lapponum*, leaf | 53 | 0 – 112 |
| | *S. lapponum*, wood | 393 | 0 – 1157 |
| | *M. trifoliata* | 81 | 48 – 125 |
| | *B. nana*, shoot | 48 | - |
| | *B. nana*, stem | 240 | - |
| | | Mean | Range |
| ***mmoX***[2] | *S. lapponum*, leaf | 7 | 4 – 12 |
| | *S. lapponum*, wood | 233 | 67 – 372 |
| | *M. trifoliata* | 30 | 19 – 37 |
| | *B. nana*, shoot | 2743 | - |
| | *B. nana*, stem | 10782 | - |

[1] Includes reads for the *amoA* genes (See Figs. 7, 8 and S6; Siljanen *et al.*, 2022)

[2] Includes reads for the non-methane mono-oxygenase genes (See Figs. 7,8 and S7; Siljanen *et al.*, 2022)

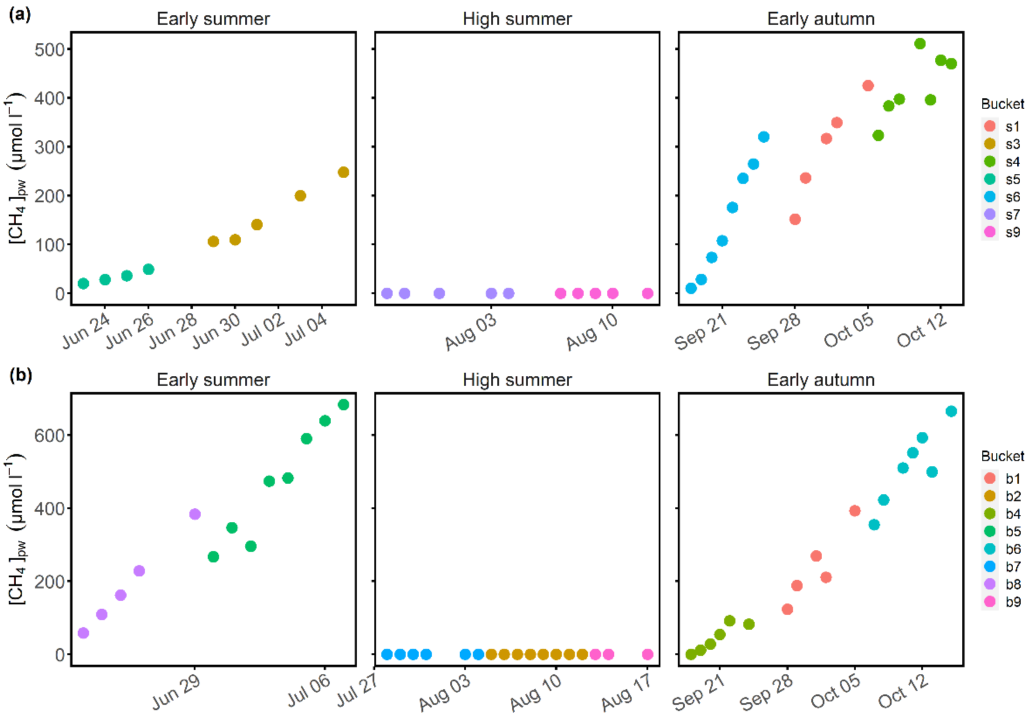


**Fig. S1** Porewater $CH_4$ concentration ($[CH_4]_{pw}$, unit: µmol $l^{-1}$) of *C. rostrata* (a) and *B. nana* (b) buckets measured in 2020. Note the different y-axis scales.

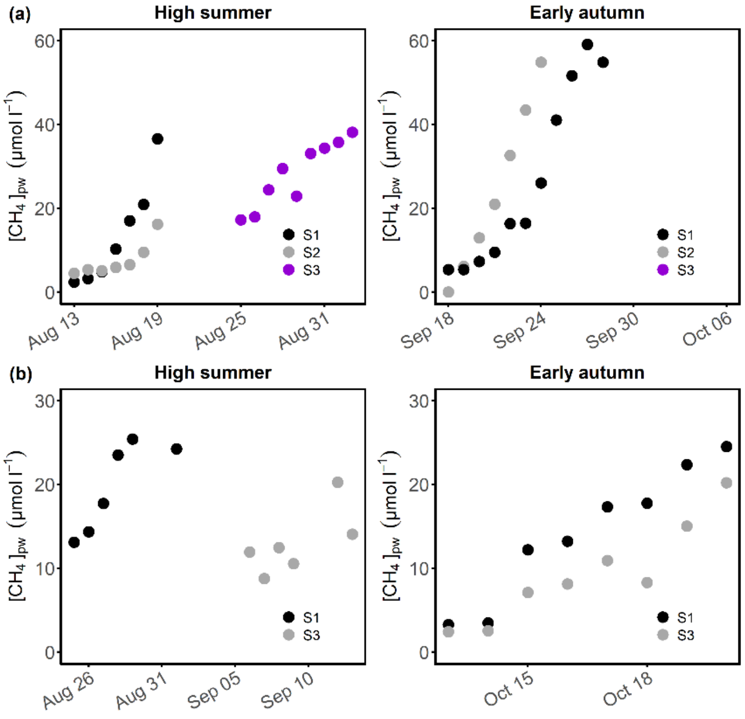


**Fig. S2** Porewater $CH_4$ concentration ($[CH_4]_{pw}$, unit: µmol $l^{-1}$) of *M. trifoliata* (a) and *S. lapponum* (b) buckets measured in 2021. No data available in early summer due GC malfunctions. Note the different y-axis scales.

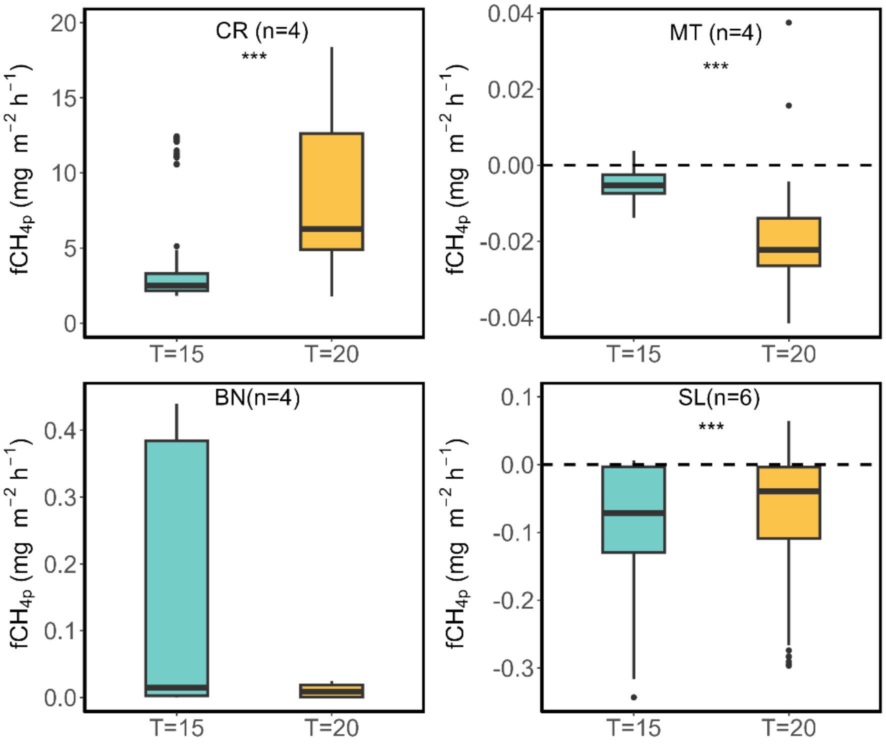


**Figure S4** Temperature effects on $CH_4$ flux ($fCH_{4p}$, mg $CH_4$ $m^{-2}$ leaf area $h^{-1}$) of *C. rostrata*, *M. trifoliata*, shrubs *B. nana* and *S. lapponum*. The number (n) of total specimens per species is less than 9 because of system malfunctions.

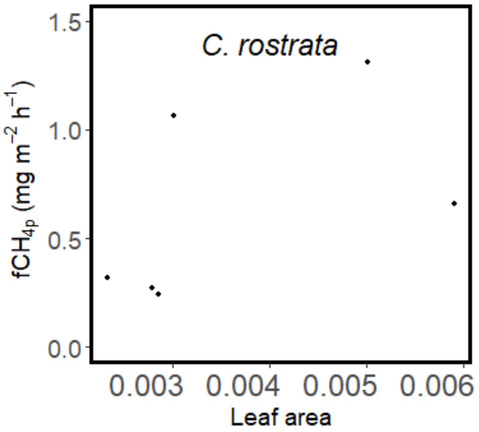


**Fig. S9** Correlation between leaf area and $CH_4$ fluxes through *C. rostrata*.

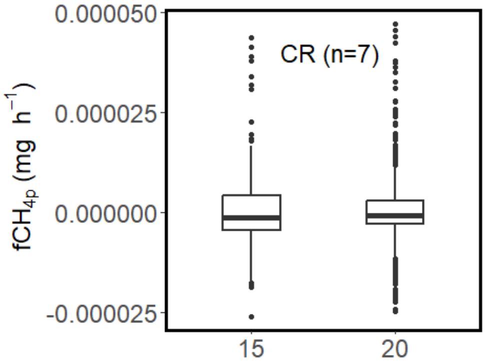


**Fig. S10** Net ecosystem exchange of *C. rostrata* (CR) before and after the increase of temperature.
